# Ubiquity and quantitative significance of bacterioplankton lineages inhabiting the oxygenated hypolimnion of deep freshwater lakes

**DOI:** 10.1101/088864

**Authors:** Yusuke Okazaki, Shohei Fujinaga, Atsushi Tanaka, Ayato Kohzu, Hideo Oyagi, Shin-ichi Nakano

## Abstract

Freshwater bacterioplankton in the oxygenated hypolimnion are reportedly dominated by specific members that are distinct from those in the epilimnion. However, no consensus exists regarding the ubiquity and abundance of these bacterioplankton, which is necessary to evaluate their ecological importance. The present study investigated the bacterioplankton community in the oxygenated hypolimnia of 10 deep freshwater lakes. Despite the broad geochemical characteristics of the lakes, 16S rRNA gene sequencing demonstrated that many predominant lineages in the hypolimnion were shared by several lakes and consisted of members occurring in the entire water layer and members specific to the hypolimnion. Catalyzed reporter deposition fluorescence *in situ* hybridization (CARD-FISH) revealed that representative hypolimnion-specific lineages, CL500–11 (*Chloroflexi*), CL500–3, CL500–37, CL500–15 (*Planctomycetes*), and the MGI group (*Thaumarchaeota*), together accounted for 1.5–32.9% of all bacterioplankton in the hypolimnion of the lakes. Furthermore, an analysis of micro-diversification based on single-nucleotide variation in the partial 16S rRNA gene sequence (oligotyping) suggested the presence of hypolimnion-specific ecotypes among the lineages occurring in the entire water layer (e.g., acI and *Limnohabitans*). Collectively, these results demonstrate the uniqueness, ubiquity, and quantitative significance of bacterioplankton in the oxygenated hypolimnion, motivating future studies to focus on their eco-physiological characteristics.

## INTRODUCTION

The hypolimnion is the dark cold water layer that lies below the thermocline in a thermally stratified lake. In a deep holomictic oligo-mesotrophic lake, this layer remains oxygenated throughout the year, as heterotrophic oxygen demand does not exceed the stock of hypolimnetic oxygen. In such a lake, accumulation and remineralization of organic matter in the hypolimnion are important biogeochemical processes (Wetzel, 2001). Recent studies have suggested that dissolved organic matter (DOM) in the oxygenated hypolimnion is enriched by the slowly consumed semi-labile fraction (Maki *et al*., 2010) and can be transformed into a more recalcitrant form by microbes (Thottathil *et al*., 2013; Hayakawa *et al*., 2016), as shown by the microbial carbon pump theory proposed for the ocean (Ogawa *et al*., 2001; Yamashita and Tanoue, 2008; Jiao and Zheng, 2011; Hansell, 2013). Other studies have demonstrated that nitrification (Small *et al*., 2013), dark carbon fixation (Callieri *et al*., 2014), and methane oxidation (Murase and Sugimoto, 2005; Bornemann *et al*., 2016) are also present in the oxygenated hypolimnion. The bacterioplankton inhabiting this realm are responsible for these important biogeochemical processes; thus, their eco-physiological characteristics should be studied.

Bacterioplankton affiliated with the phyla *Actinobacteria*, *Proteobacteria*, and *Bacteroidetes* are globally predominant in freshwater systems (Zwart *et al*., 2002; Newton *et al*., 2011). However, these data are based on surface water studies, and members of other phyla could dominate the oxygenated hypolimnion. Among them, the *Chloroflexi* CL500–11 clade (Urbach *et al*., 2001, 2007; Okazaki *et al*., 2013; Denef *et al*., 2016) and the *Thaumarchaeota* Marine Group I (MGI) group (Urbach *et al*., 2001, 2007; Auguet *et al*., 2012; Vissers *et al*., 2013; Coci *et al*., 2015; Mukherjee *et al*., 2016) are the most investigated. The relatively large cell size (1–2 μm) and high abundance (>15% of all prokaryotes) of CL500–11 suggest their quantitative importance in the oxygenated hypolimnion. MGI are ammonia-oxidizing archaea, and members in the oxygenated hypolimnion are affiliated with either *Nitrosopumilus* or *Candidatus* Nitrosoarchaeum (Coci *et al*., 2015). Other nitrifiers such as *Nitrosospira* and *Nitrospira* have also been found in the water column of the oxygenated hypolimnion (Mukherjee *et al*., 2016; Okazaki and Nakano, 2016; Fujimoto *et al*., 2016). Moreover, high-throughput sequencing of the 16S rRNA partial amplicon has highlighted members of *Planctomycetes* (e.g., CL500–3, CL500–15, and CL500–37) as representative lineages in the oxygenated hypolimnion (Rozmarynowycz, 2014; Okazaki and Nakano, 2016). These inhabitants are potentially important components of the microbial food web and biogeochemical cycling in the deep pelagic freshwater ecosystem. However, it remains unknown how ubiquitously and abundantly they are distributed in the oxygenated hypolimnion. Due to a lack of quantitative data, their ecological importance remains poorly understood.

The present study investigated the bacterioplankton community in the oxygenated hypolimnia of 10 deep freshwater lakes with a variety of geochemical characteristics. Community composition was investigated by 16S rRNA amplicon sequencing, and several representative members were microscopically characterized and quantified by CARD-FISH. This approach allowed us to create a general overview of the bacterioplankton community inhabiting the oxygenated hypolimnion of deep freshwater lakes and identify abundant and ubiquitous lineages. Moreover, analyses of their habitat preference and micro-diversification (oligotyping) facilitated hypotheses about their eco-physiological characteristics and potential diversified ecotypes.

## MATERIALS AND METHODS

### Field sampling

Water samples were collected at pelagic sites in 10 deep freshwater lakes in Japan from August to December in 2015, including Lake Mashu, Kusharo, Toya, Inawashiro, Chuzenji, Sai, Motosu, Biwa, Ikeda, and T-Reservoir (hereafter, MA, KU, TO, IN, CH, SA, MO, BI, IK, and TR, respectively) (Fig. 1). The profiles of the sampling locations are summarized in Table 1. In all, 3 to 13 depths were sampled in each lake, and the temperature and dissolved oxygen vertical profiles were measured using a CTD probe *in situ*. These lakes have permanently oxygenated hypolimnia and were thermally stratified when the sampling was carried out (Fig. 1). Total prokaryotic abundance was determined by microscopic enumeration of DAPI-stained cells (Porter and Feig, 1980).

**Fig. 1.**
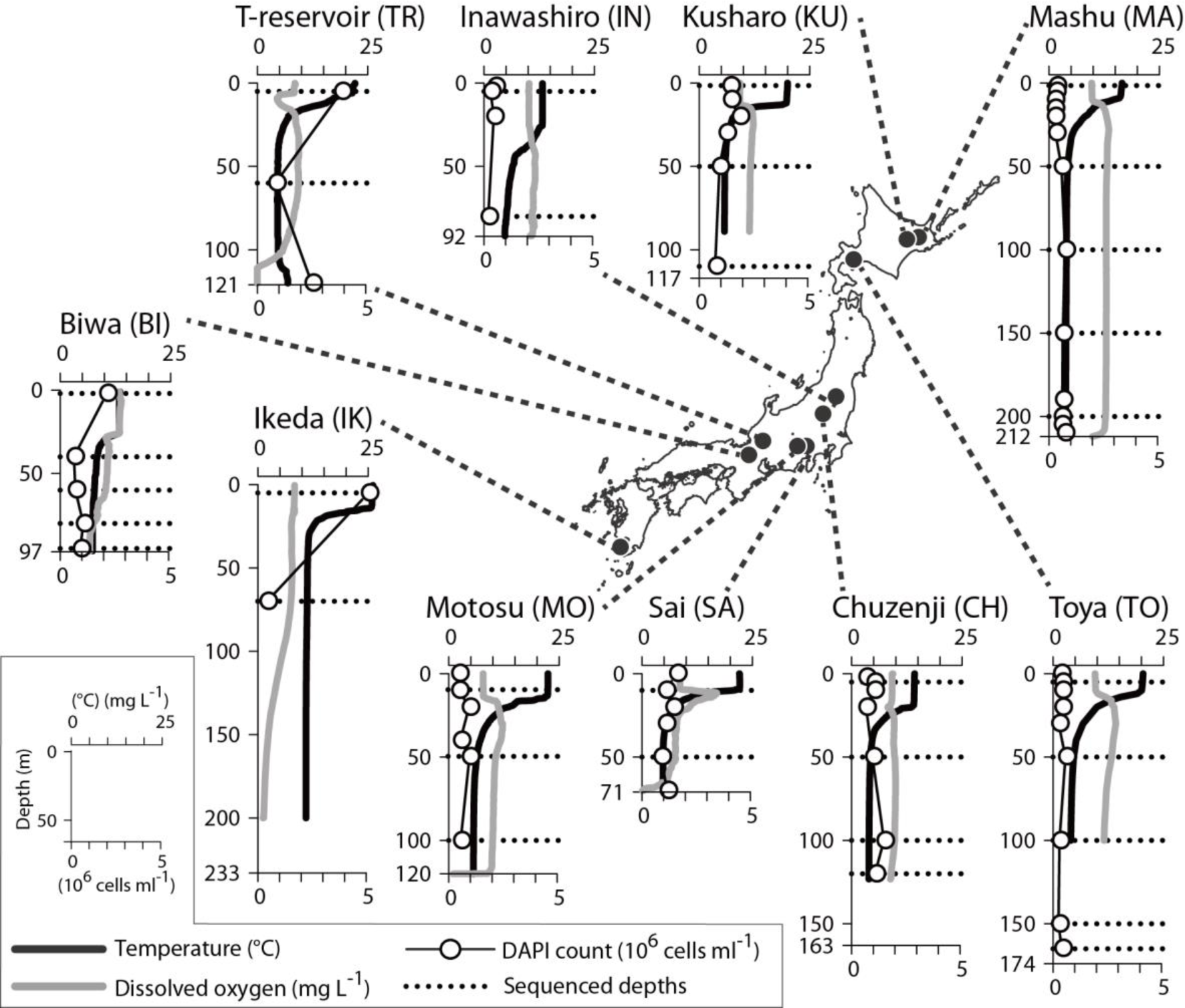
Locations and vertical profiles of the sampling sites.

**Table 1.**
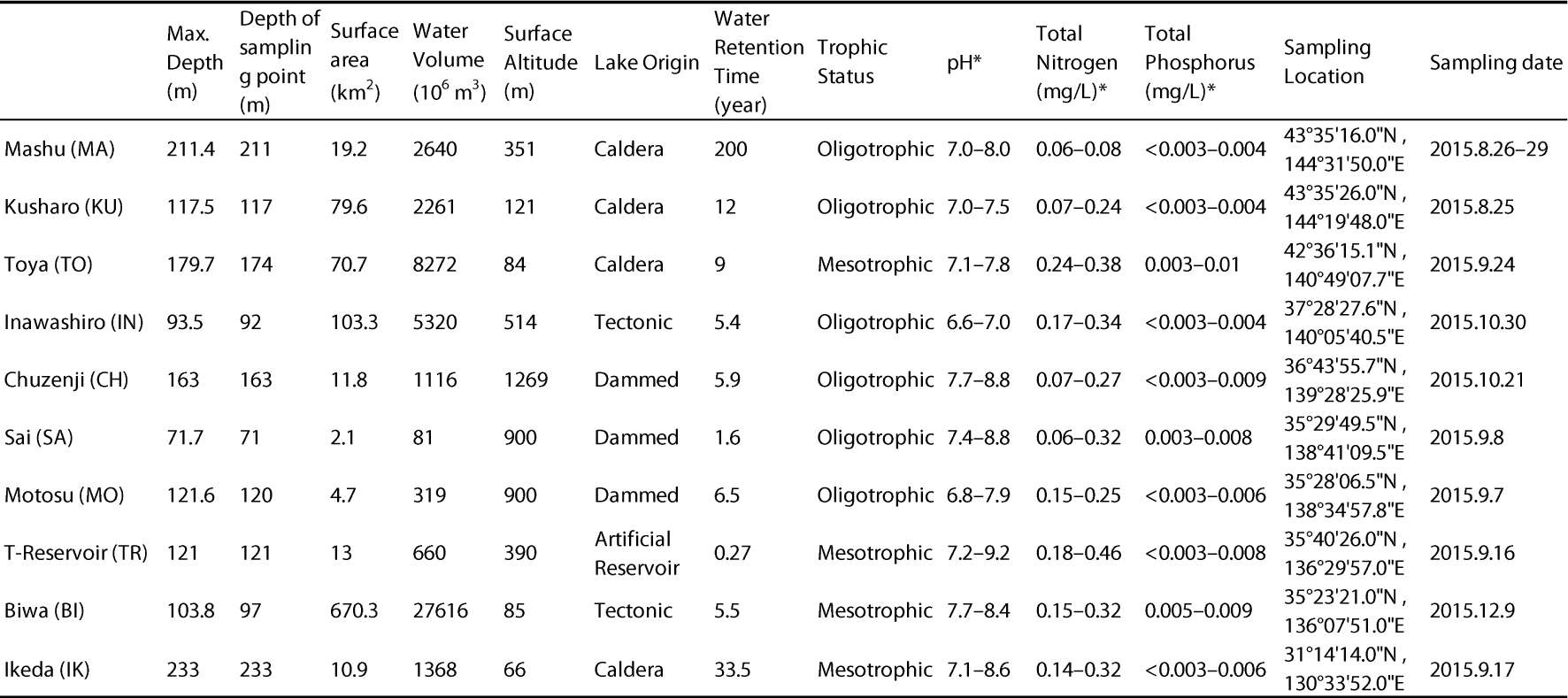
Profiles of the sites studied. Data were collected from Mori and Sato (2015) and the Japanese Ministry of the Environment public water database. *Annual range recorded in the epilimnion.

### Partial 16S rRNA gene amplicon sequencing

In total, 33 DNA samples were collected (Fig. 1). Immediately after sampling, 70–300 mL the water collected was filtered through a 0.2 ¼m polycarbonate filter (47 mm diameter; Whatman, Maidstone, UK). The filter was stored at -20°C until DNA was extracted using the PowerSoil DNA Isolation Kit (MoBio Laboratories, Carlsbad, CA, USA). The V4 and V5 regions of the 16S rRNA gene were amplified using the modified 530F and 907R primers (Nunoura *et al*., 2009) and then an eight-base-pair DNA tag (for post-sequencing sample identification) and 454 adaptors were conjugated by second PCR, as described previously (Okazaki and Nakano, 2016). The PCR products from the samples were pooled in equimolar quantities and sequenced in the 1/8 regions of a sequencing reaction on the Roche 454 GS-FLX Titanium sequencer (Roche Science, Mannheim, Germany). The nucleotide sequence data are available in the Sequence Read Archive database under accession numbers DRX062810–DRX062842 (BioProject: PRJDB5151).

### Analysis of sequencing reads

The sequence data were processed using the UPARSE pipeline (Edgar, 2013) following the author’s instructions (http://www.drive5.com/usearch/manual/upp_454.html). We used a fastq_maxee value of 1.0, a truncated length of 350 bp, and an OTU creation identity threshold of 97%. Thereafter, respective OTUs were taxonomically assigned with the SINA 1.2.11 (Pruesse *et al*., 2012) online tool (https://www.arb-silva.de/aligner/) referring to the 123 Ref database (Quast *et al*., 2013) and using the default parameter settings. Subsequently, non-prokaryotic OTUs (i.e., chloroplast, eukaryote, and unclassified domains) were removed. The resulting 96 149 reads were used for subsequent analyses.

Before calculating community diversity, coverage-based rarefaction (Chao and Jost, 2012) was applied. Reads were randomly discarded from each sample until coverage was <96.59% (i.e., slope of the rarefaction curve was >0.0341), which was the minimum value recorded among samples. Subsequently, alpha (Shannon diversity) and beta diversities (non-metric multidimensional scaling [NMDS] based on Bray–Curtis dissimilarity) were calculated. These analyses were performed using the vegan 2.4–0 package (Oksanen *et al*., 2016) in R 3.3.1 software (http://www.R-project.org/).

The OTUs were classified following the nomenclature proposed by Newton *et al*. (2011). Using the “‐‐search” option in the SINA 1.2.11 (Pruesse *et al*., 2012) stand-alone tool, the representative sequences of the respective OTUs (generated by the UPARSE pipeline) were searched and classified against the original ARB (Ludwig *et al*., 2004) database provided by Newton *et al*. (2011). Dominant OTUs that failed to be classified by this procedure were named manually. If closely related (>99% identity) sequences with the prefixes “CL” (Urbach *et al*., 2001) or “LiUU-” (Eiler and Bertilsson, 2004) were in the public sequence database, their names were preferentially used. The *Planctomycetes* phylogenetic clades were newly defined for the collective descriptions (Fig. S1). In other cases, the OTU was shown by its taxonomic affiliation (e.g., genus) based on the SILVA nomenclature.

The sequences were aligned using the SINA 1.2.11 (Pruesse *et al*., 2012) stand-alone tool against the SILVA 123 Ref NR 99 database to construct a phylogenetic tree (Quast *et al*., 2013). Maximum Likelihood trees were calculated by the RAxML 8.2.4 software (Stamatakis, 2014) with the GTR substitution model and the GAMMA rate heterogeneity model. The trees were drawn in MEGA 7 software (Kumar *et al*., 2016).

### Oligotyping

Intra-OTU micro-diversity of representative lineages was analyzed by oligotyping, which facilitated detection of single nucleotide variation by excluding the effects of sequencing errors based on the Shannon entropy values (Eren *et al*., 2013). Following the author’s instructions (http://merenlab.org/software/oligotyping/), the quality filtered FASTA file generated by the UPARSE pipeline was split at individual OTUs and aligned against the Greengenes (DeSantis *et al*., 2006) alignment database using the SINA 1.2.11 (Pruesse *et al*., 2012) stand-alone tool. Uninformative columns were removed by subsequently applying the “o-trim-uninformative-columns-from-alignment” and “o-smart-trim” scripts. Several rounds of oligotyping were repeated by manually choosing the most informative (i.e., the highest entropy) column until all oligotypes with > 100 reads exceeded the purity score of 0.90. The minimum substantive abundance parameter (option-M) was always set to 10.

### CARD-FISH

CARD-FISH was performed based on Pernthaler *et al*. (2002) with some modifications as described previously (Okazaki *et al*., 2013). Specifically, the hybridization, amplification, and washing steps were carried out at 46°C. AlexaFluor 488 (Life Technologies, Carlsbad, CA, USA) was used as the tyramide-conjugated fluorescent dye. The oligonucleotide probes were designed previously for CL500–11 (Okazaki *et al*., 2013) and MGI (Coci *et al*., 2015), and newly designed for the other targets (Table 2). The probes were constructed using the “Design Probes” function in ARB 6.0.3 software (Ludwig *et al*., 2004) against the SILVA 123 Ref NR 99 database (Quast *et al*., 2013). Specificity of the probes was confirmed by an NCBI BLAST online search and the Test Probe 3.0 tool against the SILVA 123 Parc database (https://www.arb-silva.de/search/testprobe/). The probes designed in this study targeted the V4 or V5 regions of the 16S rRNA gene, which were read by amplicon sequencing. We confirmed that the probes perfectly and exclusively matched all sequencing reads affiliated with the target lineage. To enhance the fluorescent signal, oligonucleotide helpers (Fuchs *et al*., 2000) were used for several probes (Table 2). The helpers were designed to target the adjacent or opposite loci of the probe target site to loosen the secondary rRNA structure (Fuchs *et al*., 2000) by confirming that all probe-targets in the database were not mismatched with their corresponding helpers. The hybridization buffer contained 0.5 μg mL-^1^ probe and 0.1 μg mL-^1^ of each helper. The formamide concentration in the buffer was determined for each probe (Table 2) by testing a series of concentrations to obtain the best stringency (i.e., highest concentration without signal loss). The stringency test was performed in samples with a high read proportion of the target determined by amplicon sequencing. The hybridized filters were counterstained with DAPI and enumerated under an epifluorescence microscope. At least 300 DAPI-positive cells and the corresponding FISH-positive cells were enumerated three times per sample (the same filter piece). A negative NON338 probe control (Wallner *et al*., 1993) confirmed that no false-positive cells were present.

**Table 2.**
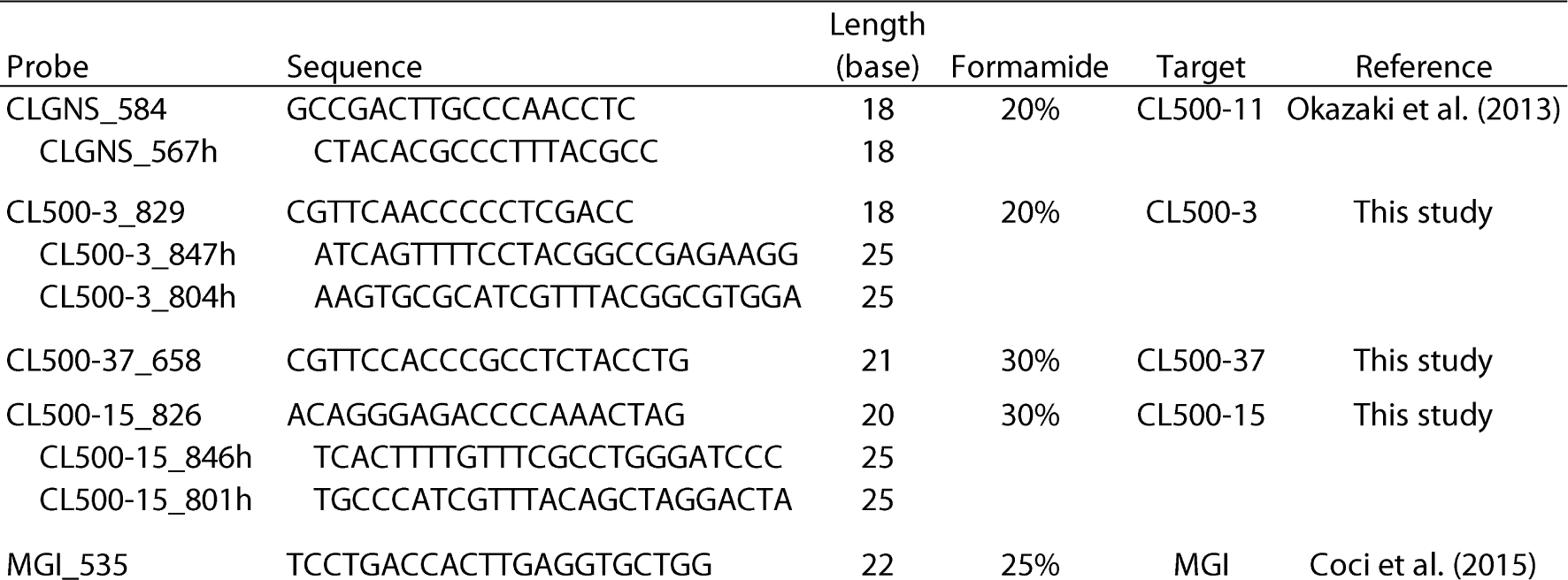
Catalyzed reporter deposition fluorescence *in situ* hybridization (CARD-FISH) probes, helpers, and formamide concentrations (at 46°C) used in this study.

## RESULTS

16S rRNA partial amplicon sequencing generated 666 OTUs from the 96 149 reads originating from the 33 samples collected from the water columns of the 10 lakes. The complete dataset generated by the pipeline is available in Supplementary Information.

Phyla *Actinobacteria*, *Bacteriodetes*, and class *Betaproteobacteria* generally dominated throughout the water column, whereas *Chloroflexi* and *Planctomycetes* were also predominant phyla in the hypolimnion (Fig. 2). The epilimnetic and hypolimnetic communities were compared at the OTU level by averaging the samples from each layer (Fig. 2). The dominant members in the epilimnion were generally shared between the lakes (e.g., acI-B1, acI-A7, Lhab-A1, and bacI-A1). In addition to the lineages common to the epilimnion, *Chloroflexi* and *Planctomycetes* were also ranked as dominant OTUs in the hypolimnion (Fig. 2). CL500–11 alone accounted for most of the *Chloroflexi* reads, whereas *Planctomycetes* consisted of diverse OTUs (e.g., CL500–3, CL500–15, CL500–37, and plaI-A) (Fig. 2), which were affiliated with three classes in the phylum (Fig. S1). The alpha (Shannon) diversity was higher in the hypolimnion than that in the epilimnion of all 10 lakes (Fig. 2). The beta diversity analysis (NMDS) clearly separated the hypolimnetic communities from the epilimnetic communities, but no clear separation between the lakes was observed (Fig. 3).

**Fig. 2.**
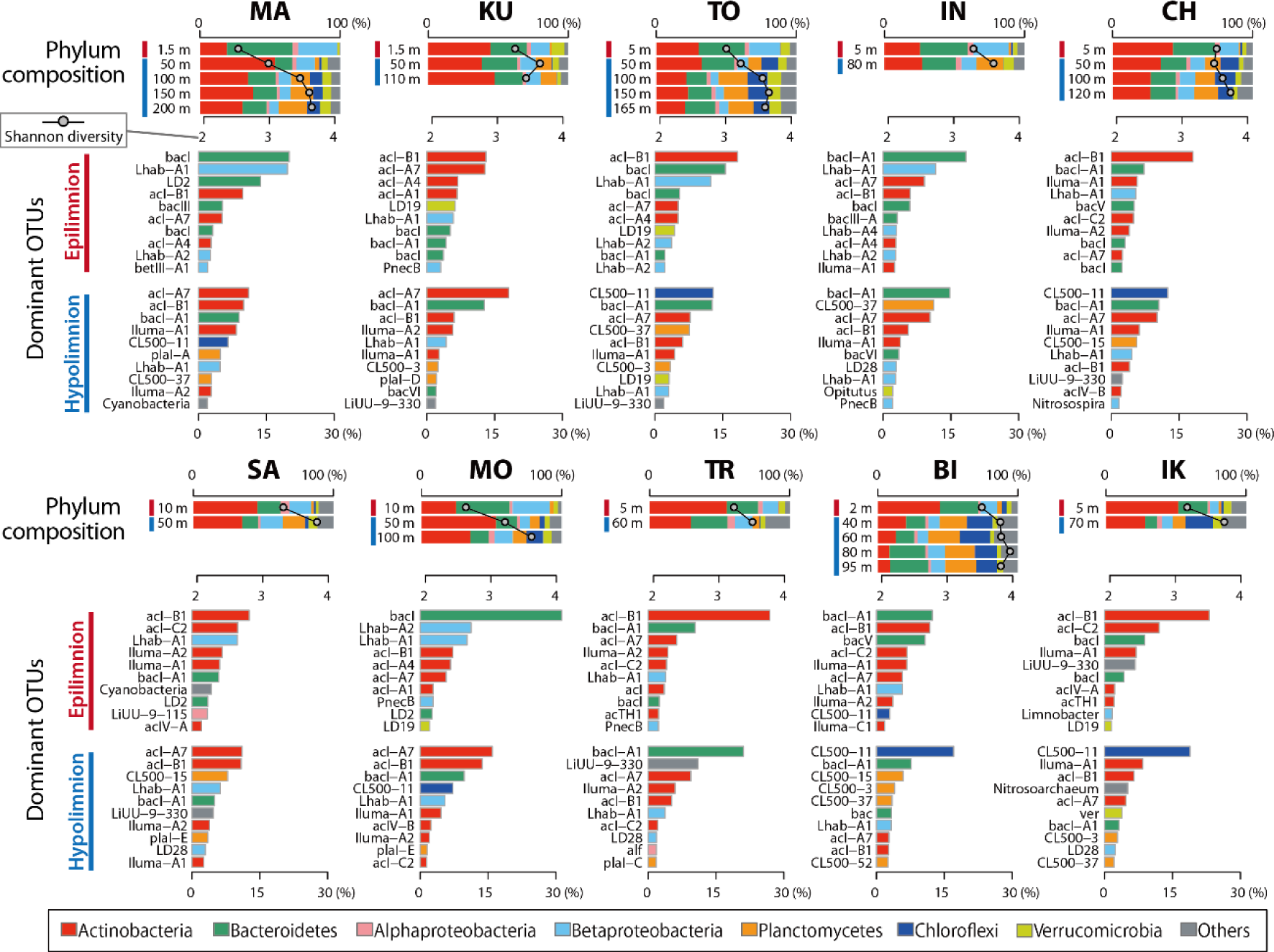
Composition of the 16S amplicon reads. The top panel (band graphs) displays phylum-resolved community composition at each depth for each lake, with an overwritten line graph indicating the alpha (Shannon) diversity. The two lower panels show the 10 dominant operational taxonomic units (OTUs) in the epilimnion and hypolimnion, composed of averaged data for each layer (the depths averaged are illustrated by red and blue lines in the top panel). Bar colors indicate phyla to which individual OTUs were assigned.

**Fig. 3.**
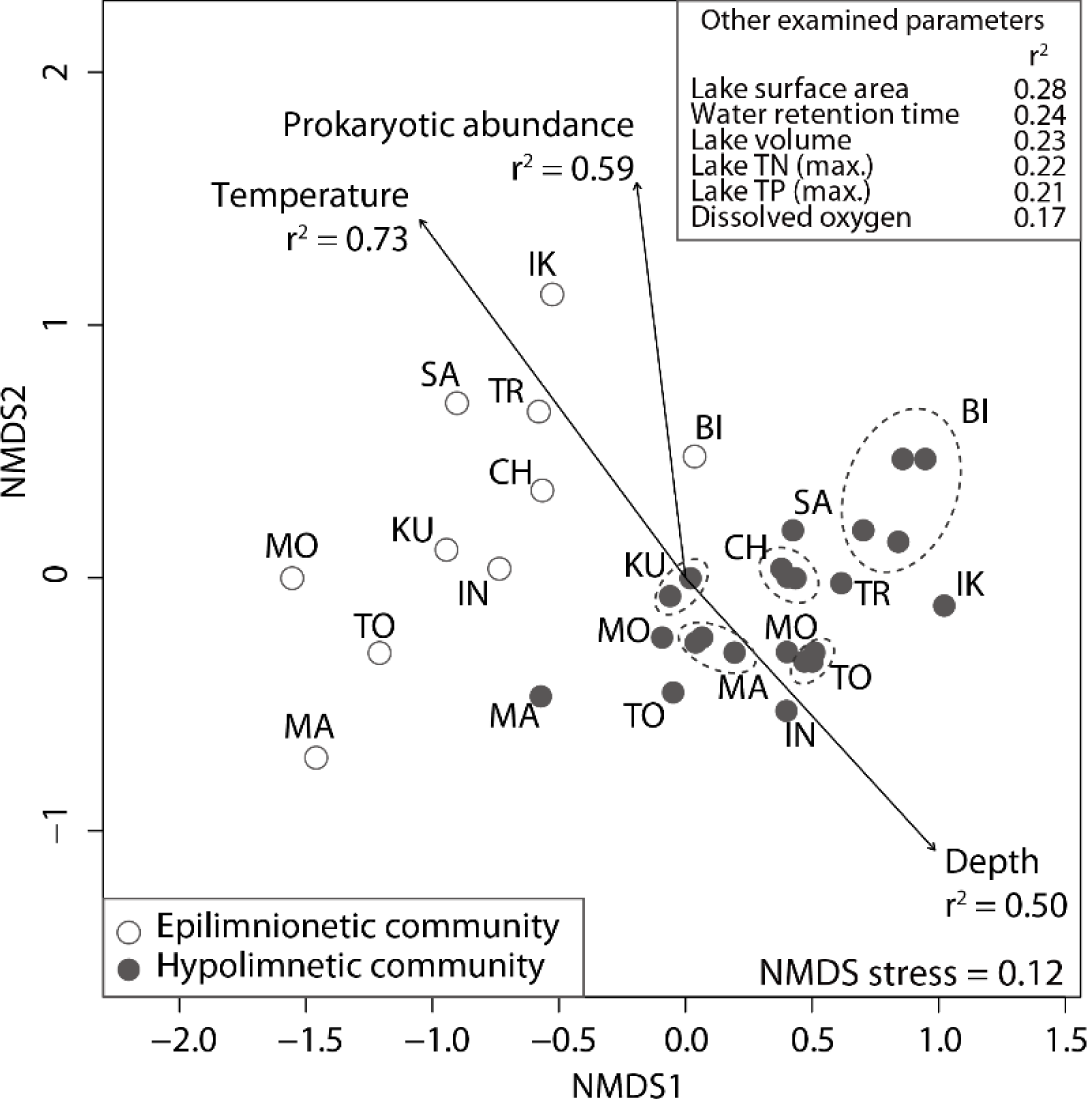
Non-metric multidimensional scaling (NMDS) ordination of all 33 bacterioplankton communities. The arrows are fitted vectors for the environmental variables calculated using the envfit function. The direction of the arrow indicates the direction at which the gradient of the environmental variable was maximum. The length of the arrow is proportional to the squared correlation coefficient (r2). Only significant variables (p < 0.01; based on 999 permutations) were visualized. The other variables examined are shown in the top right box with the r^2^ values.

Vertical preference of the bacterioplankton in each OTU was determined based on the number of lakes where it was abundant (>1% of all amplicon reads) in each water layer (Fig. 4). Seven OTUs were defined as “whole-layer generalists,” which were abundant in either the epilimnion or hypolimnion of more than eight lakes (Fig. 4). Twenty-two OTUs were defined as “hypolimnion specialists,” which were abundant in the hypolimnion of multiple lakes but not abundant in the epilimnion of almost all (nine or ten) lakes. Conversely, 19 OTUs were identified as “epilimnion specialists” (Fig. 4). Several bacterioplankton in the hypolimnion showed a habitat preference between the lakes, but only few environmental parameters significantly explained the patterns (Fig. 5). The oligotyping analysis revealed that the whole-layer generalists exhibited more intra-OTU diversity than that of the hypolimnion specialists (Fig. 5). The composition of the oligotypes in each sample is shown in Supplementary Figure S2.

**Fig. 4.**
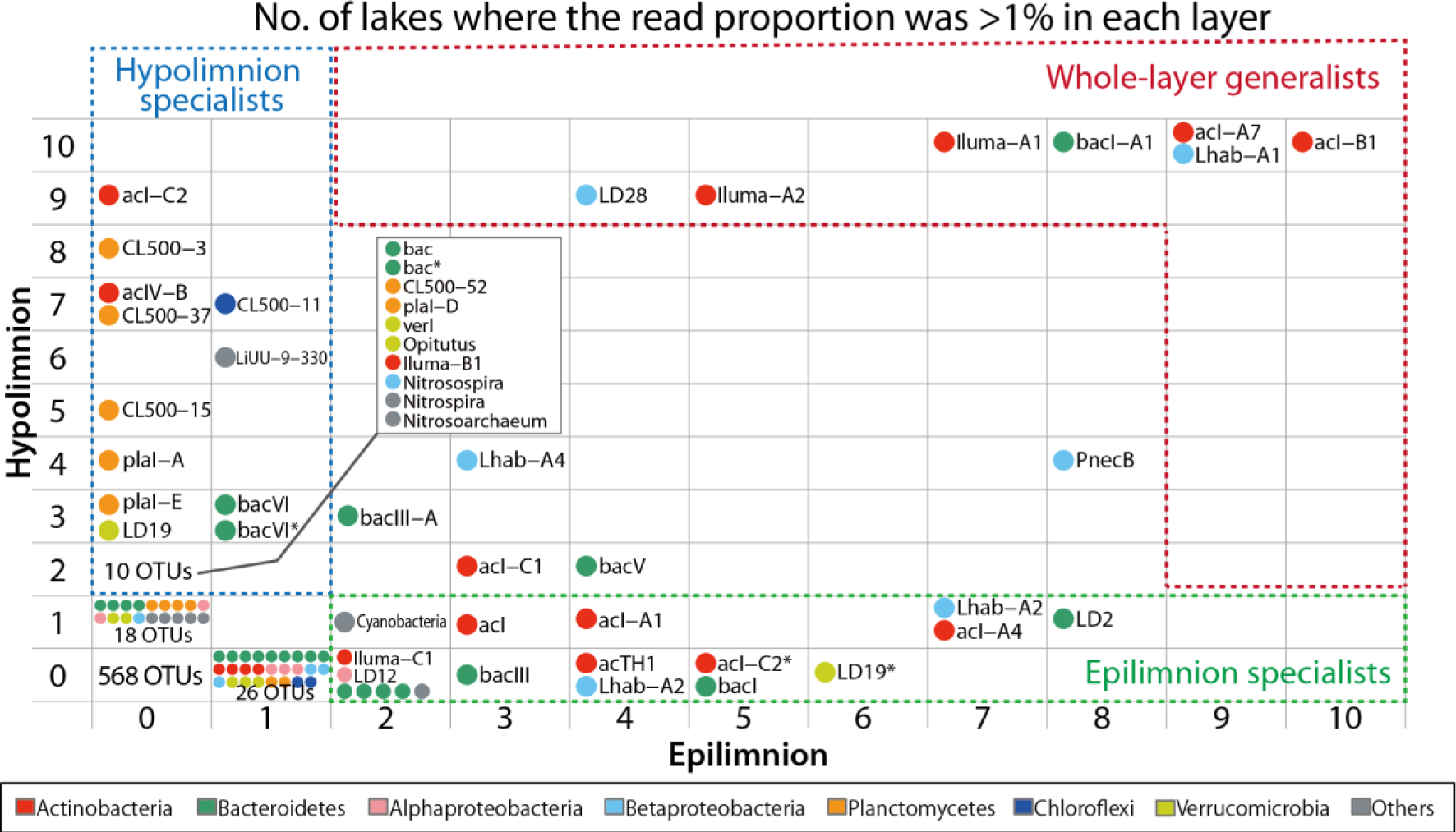
Vertical preferences of individual operational taxonomic units (OTUs), mapped by the number of lakes where individual OTUs accounted for >1% of all amplicon reads in each water layer. Data for the hypolimnion were generated by averaging the data at multiple depths in the hypolimnion. Dotted boxes highlight the groups defined in this study (see main text). Point color indicates the phylum to which an individual OTU was assigned. Asterisks in the group name distinguish the different OTUs assigned to the same group.

The newly constructed FISH probes targeted a monophyletic clade of the target lineages (Fig. S3). Enumeration of the positive cells revealed that CL500–11 accounted for >10% of all prokaryotic cells in four lakes (maximum was 25.9% at 60 m in BI). CL500–3, CL500–37, and MG1 respectively accounted for >3% in several lakes. CL500–15 were less abundant but still detectable, with a maximum percentage of 1.6% (Fig. 6). The cells detected in each target shared identical morphology between the lakes: CL500–11 were curved rods 1–2 μm long; CL500–3, CL500–37, and CL500–15 were cocci approximately 1 μm diameter; and MG1 were rods around 1 μm long (Fig. 6). CL500–15 cells were often found to be particle-associated (51% of the cells detected at 50 m in SA), whereas the other lineages were mostly planktonic. The correlation between relative abundance determined by amplicon sequencing and CARD-FISH was significant for all targets, but the CARD-FISH estimates tended to be lower than those of amplicon sequencing (Fig. S4).

## DISCUSSION

### Structure of the bacterioplankton community in the oxygenated hypolimnion

The present study investigated lakes with a variety of environmental characteristics, ranging from a cold (hypolimnetic temperature = 4°C) oligotrophic lake (MA) to a warm (11°C) mesotrophic lake (IK) (Fig. 1 and Table 1). Nevertheless, the bacterioplankton communities in the oxygenated hypolimnion were relatively similar among the lakes and separated from communities in the epilimnion (Fig. 3), indicating that the oxygenated hypolimnion is an independent niche with distinct microbial communities. The community consisted of members that were ubiquitous and predominant across the lakes and water layers (whole-layer generalists), and members that exclusively inhabited the oxygenated hypolimnion (hypolimnion specialists) (Fig. 4). The whole-layer generalists were composed of several commonly known freshwater bacterioplankton lineages (e.g., acI, Iluma-A1, Iluma-A2, Lhab-A1, LD28, and bacI-A1), whereas the hypolimnion specialists were represented by phyla that were not common to the epilimnion, including *Chloroflexi* CL500–11, members of *Planctomycetes* (e.g., CL500–3, CL500–15, CL500–37, and plaI-A), and *Ca*. Nitrosoarchaeum in *Thaumarchaeota* (Figs. 2, 4). Their abundance and ubiquity were demonstrated by the CARD-FISH analysis, showing that they collectively accounted for 1.5% (TR) to 32.9% (BI) of all bacterioplankton in the hypolimnion. Our analysis also revealed that these groups were not cosmopolitan and were actually absent in some lakes (e.g., IN and KU for CL500–11, CH for CL500–3, and MO for CL500–37) (Figs. 5, 6), suggesting that they have respective habitat preferences and ecological niches.

**Fig. 5.**
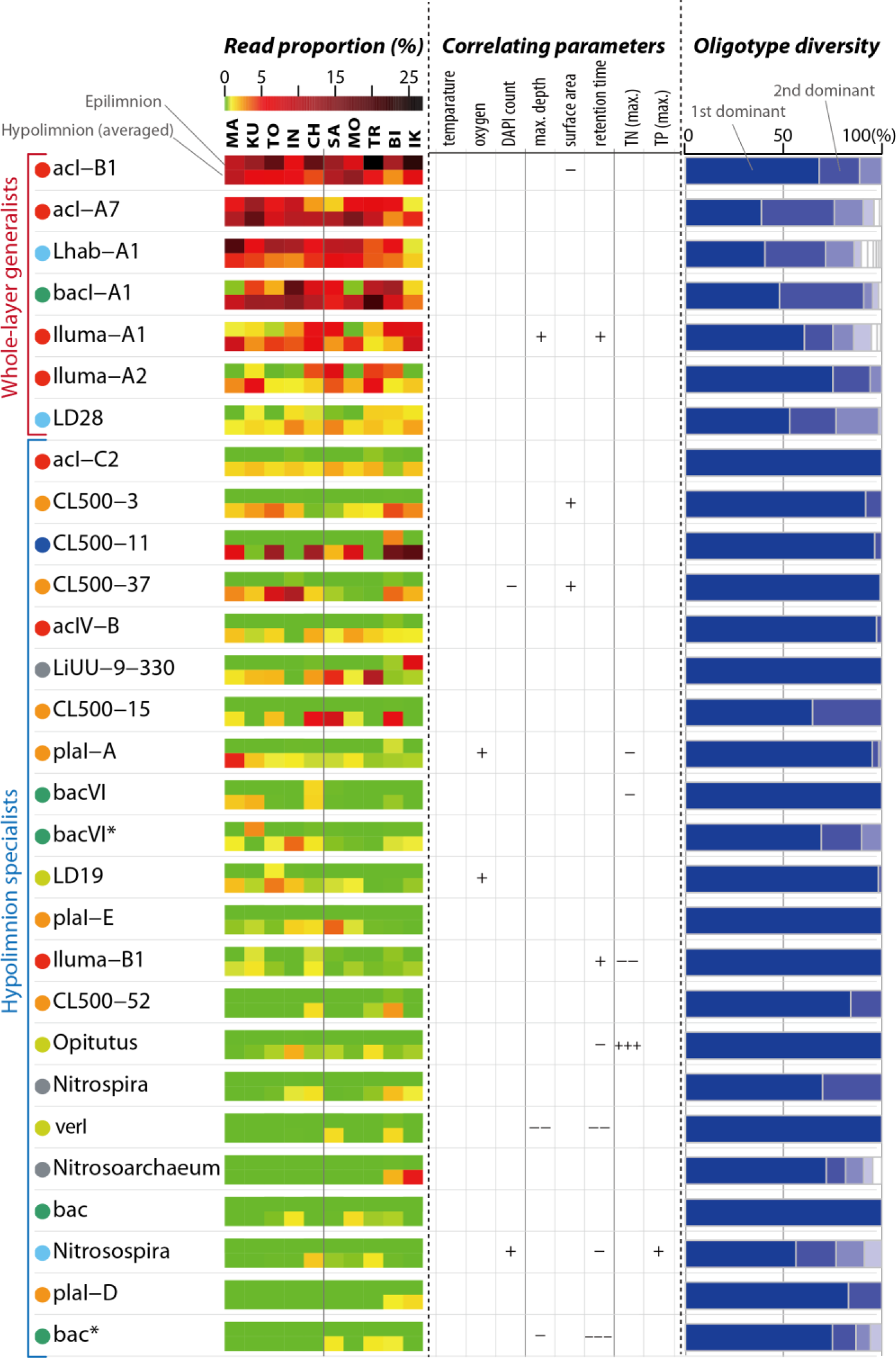
Distribution patterns (left column), correlating environmental parameters (center), and oligotype diversity (right) of the hypolimnion inhabitants. The distribution pattern is illustrated by the read proportion to total amplicon reads. The hypolimnion data were generated by averaging the data at multiple depths in the hypolimnion. The correlation between read proportion and environmental parameters in the hypolimnion was evaluated by Spearman’s test. For a positive correlation, +++, p < 0.005; ++, p < 0.01; +, p < 0.05. For a negative correlation, “-” was shown instead of “+”. Maximum values recorded in TN and TP were used based on the database. The right column (band graphs) indicates composition of the oligotypes among all amplicon reads assigned to each operational taxonomic unit (OTU). Point colors indicate phyla to which individual OTUs were assigned (See Figs. 2 and 4 for legend). Asterisks in the group name distinguish the different OTUs assigned to the same group.

### Ubiquity, quantitative importance, and potential eco-physiology of the hypolimnion specialists

The predominance (~25% of all bacterioplankton) of planktonic CL500–11 cells in several lakes (Fig. 6) suggests that their resources are diffuse, abundant and ubiquitous. The metagenome-assembled genome and *in situ* transcriptional evidence of CL500–11 in Lake Michigan suggests their importance in peptide turnover (Denef *et al*., 2016). Peptides in aquatic systems are mainly derived from peptidoglycans in the bacterial cell wall (McCarthy *et al*., 2013; Nagata *et al*., 2003) and from proteins released by other bacteria (Tanoue *et al*., 1995) or phytoplankton (Nguyen and Harvey, 1997; Yamada *et al*., 2012). Previous studies in BI have collectively demonstrated that N-rich (by stoichiometry) or protein-like (by fluorescence properties) semi-labile dissolved organic matter (DOM) derived from autochthonous phytoplankton production that accumulates in the hypolimnion is slowly remineralized during stratification (Kim *et al*., 2006; Maki *et al*., 2010; Thottathil *et al*., 2013). Thus, it is possible that CL500–11 is scavenging protein-like debris that accumulates in the lake due to its relatively recalcitrant nature. Given that the water retention time of a system affects DOM composition (Kellerman *et al*., 2014; Catalán *et al*., 2016) and that autochthonous dissolved proteins can accumulate in a lake even at centennial time scales (Goldberg *et al*., 2015), we expected that lakes with a longer water retention time would contain more bacterioplankton lineages specialized to consume relatively recalcitrant DOM. In the present study, the water retention time of the lakes ranged from 0.27 years (TR) to 200 years (MA) (Table 1). However, most of the hypolimnion specialists, including CL500–11, were not distributed in a manner that was associated with water retention time (Fig. 5). Nonetheless, it is still possible that a water retention time (<2 years) too short to accumulate semi-labile DOM would have resulted in the absence of CL500–11 in SA and TR (Fig. 6). The absence of CL500–11 in KU and IN (Fig. 6) is more challenging to explain, but may have resulted from the low pH compared to the other lakes (Table 1). As CL500–11 is a large cell (Fig. 6), protistan size-selective grazing (Pernthaler, 2005) may also be a factor controlling CL500–11 dynamics, and little information is available on the grazer communities inhabiting the deep oxygenated hypolimnion (Masquelier *et al*., 2010; Mukherjee *et al*., 2015). Although these assumptions are speculative, future studies should verify them given their ubiquity and quantitative importance in deep freshwater systems. Indeed, the dominance of CL500–11 has been reported in the two largest deep freshwater systems on Earth, the Laurentian Great Lakes (Rozmarynowycz, 2014; Denef *et al*., 2016) and Lake Baikal (Kurilkina *et al*., 2016).

**Fig. 6.**
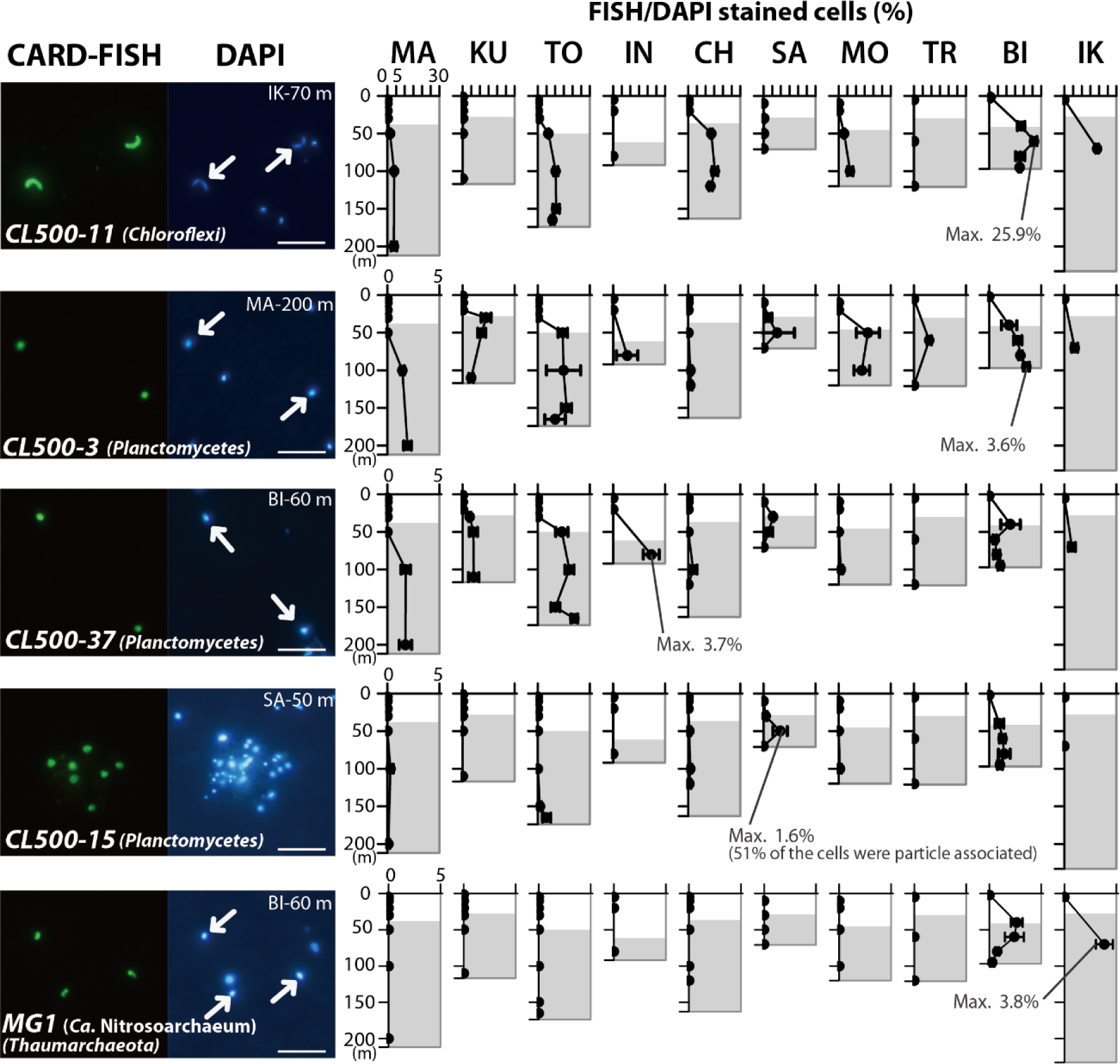
Catalyzed reporter deposition fluorescence *in situ* hybridization (CARD-FISH) images and the enumeration results. Positive cells and the corresponding DAPI-stained image are shown in each micrograph. Arrows in the DAPI-image indicate cells with FISH-positive signals. Scale bar = 5 μm. The line graphs show percentages of FISH-positive cells to DAPI-positive cells, and the error bars indicating the standard deviation determined by triplicate enumeration of an identical filter. The maximum value recorded for each lineage is designated. Gray background illustrates the hypolimnion (i.e., below the thermocline).

CL500–3 and CL500–37 were the two most abundant *Planctomycetes* in the present study (Fig. 6). Each was affiliated with their respective phylogenetic clade in the class *Phycisphaerae* (Fig. S1). Aquatic *Planctomycetes* are often associated with algal blooms (Morris *et al*., 2006; Pizzetti *et al*., 2011), and genomic evidence indicates their potential to aerobically consume sulfated polysaccharides derived from algae (Glöckner *et al*., 2003; Woebken *et al*., 2007; Erbilgin *et al*., 2014). In a marine study, sequences closely related to CL500–3 were enriched in DNA extracted from bacterioplankton that assimilate protein secreted by phytoplankton (Orsi *et al*., 2016). Consequently, it can be hypothesized that CL500–3 and CL500–37 consume polysaccharides or proteins derived from phytoplankton. Given that no photosynthesis occurs in the dark hypolimnion, these organisms presumably rely on substrates released from sinking phytoplankton. Indeed, substantial sinking fluxes of green algae (*Staurastrum*), a diatom (*Fragilaria*), and cyanobacteria (*Synechococcus*) into the hypolimnion have been reported in BI (Kagami *et al*., 2006; Takasu *et al*., 2015), where CL500–3 and CL500–37 were both abundant (Fig. 6). Their dependence on pelagic autochthonous resources was also supported by the finding that their read proportions were positively correlated with lake surface area (Fig. 5). However, they did not always co-occur, and disproportional dominance of CL500–3 was found in MO and TR, and of CL500–37 in CH and IN (Fig. 6). Amplicon sequencing data taken from the oxygenated hypolimnion in Lake Michigan (Fujimoto *et al*., 2016) showed that only CL500–37 was abundant (Fig. S5). These observations indicate that the ecological niches of CL500–3 and CL500–37 are similar but not the same. Given that algal exudates from different phytoplankton species select different bacterial communities (Šimek *et al*., 2011; Paver *et al*., 2013), the difference might be attributable to differences in the sinking phytoplankton species in a lake. It should also be noted that their closely related sequences were not necessarily retrieved from the oxygenated hypolimnion and have been found in an Antarctic lake (Karlov *et al*., 2016), arctic lake (Ntougias *et al*., 2016), and Baltic Sea ice (Eronen-Rasimus *et al*., 2015) (Fig. S3A). More information is needed to elucidate the eco-physiological characteristics of these widespread and abundant yet largely overlooked bacterial lineages.

The CL500–15 clade belonged to the uncultured OM190 class (Fig. S1) with only three sequences in the database: two from deep freshwater lakes (Urbach *et al*., 2001; Pollet *et al*., 2011) and one from deep sea sediment (Zhang *et al*., 2013) (Fig. S3B). Another sequence was reported from the littoral water of Lake Baikal (Parfenova *et al*., 2013). In the present study, CL500–15 was detected in half of the lakes (Fig. 4), indicating that they are one of the most common lineages in the oxygenated hypolimnion. The FISH analysis revealed a high proportion of particle-associated cells (Fig. 6) and microscopic observations revealed that the particles were not cells but rather looked like transparent exopolymer particles (TEP), which are gel-like sticky particles mainly composed of polysaccharides (Passow, 2002). Members of the OM190 class in marine systems have been preferentially detected in the particle-associated fraction (Salazar *et al*., 2015; Bižić-Ionescu *et al*., 2015), suggesting that their particle-associated form is preserved across the class. Particle-associated bacteria can contribute disproportionally to total bacterial activity (Lemarchand *et al*., 2006; Grossart *et al*., 2007), and CL500–15 is presumably one of the major TEP decomposers that plays a substantial role in substrate remineralization in the oxygenated hypolimnion.

Other representative *Planctomycetes* were affiliated with class *Planctomycetacia* (plaI-A–F) (Fig. S1). They usually had a smaller proportion of reads than those of CL500–3, CL500–37, and CL500–15 (Fig. 2). However, plaI-A was the most represented *Planctomycetes* based on the read proportion in the oxygenated hypolimnion of MA (Fig. 2) and Lake Michigan (Fig. S5). Each of the other members of plaI (e.g., plaI-B [CL500–52], plaI-D, and plaI-E) showed their respective distribution patterns between the lakes (Fig. 5). Overall, *Planctomycetacia* (plaI group) was generally less ubiquitous and abundant, but more diverse than the other two classes (i.e., *Phycisphaerae* and OM190) inhabiting the oxygenated hypolimnion.

In the present study, the MG1 group was detected only from two lakes (BI and IK) (Fig. 5), and their maximum percentage determined by FISH was 3.8% (Fig. 6). These numbers are lower than those of a previous study that detected 8.7–19% MG1 in the oxygenated hypolimnion of all six subalpine lakes investigated (Callieri *et al*., 2016). In the present study, MG1 was exclusively affiliated with *Ca*. Nitrosoarchaeum (Blainey *et al*., 2011), but *Nitrosopumilus*, another predominant MG1 member in the oxygenated hypolimnion (Berdjeb *et al*., 2013; Vissers *et al*., 2013; Coci *et al*., 2015), was not detected. *Ca*. Nitrosoarchaeum has been reported in the oxygenated hypolimnion of Crater Lake (Urbach *et al*., 2001, 2007), Lake Redon (Auguet *et al*., 2012), and Lake Superior (Mukherjee *et al*., 2016), which are oligotrophic lakes with a hypolimnetic temperature of 4°C. The occurrence of *Ca*. Nitrosoarchaeum in BI and IK, which are mesotrophic lakes with a hypolimnetic temperature of 8–11°C (Fig. 1), revealed their broad habitat spectrum. In other lakes (e.g., CH and TR), ammonia-oxidizing bacteria, *Nitrosospira*, were detected in the oxygenated hypolimnion (Fig. 5), in agreement with the previously suggested niche separation between ammonia-oxidizing archaea and bacteria (Coci *et al*., 2015; Mukherjee *et al*., 2016). Another nitrifier, *Nitrospira*, was also detected in the oxygenated hypolimnion (Fig. 5), in line with previous studies (Small *et al*., 2013; Okazaki and Nakano, 2016). However, no nitrifiers were detected in the three northern lakes (MA, KU, and TY) (Fig. 5). Notably, their absence should not be concluded by the present data, as the nitrifier community can shift over seasons (Okazaki and Nakano, 2016). Nevertheless, our results indicate the potential diversity of the nitrification systems in the oxygenated hypolimnion, yet information remains scarce to conclude the cause and effects of the diversity.

The discussion above is based on the assumption that each of the bacterioplankton lineages prefer their particular suitable habitat. However, it is also possible that the occurrence of the members is controlled by occurrence of other lineages, as bacterioplankton often have streamlined genomes and are dependent on each other for lost metabolic functions (Morris *et al*., 2012; Garcia *et al*., 2015; Mas *et al*., 2016). In the present study, many pairs of hypolimnion specialists showed correlating distribution patterns between the lakes (Fig. S6). For example, CL500–3 was positively correlated with CL500–37 and *Ca*. Nitrosoarchaeum, and negatively correlated with *Nitrosospira* (Fig. S6). Although these results do not directly support an interaction, they suggest that some hypolimnion specialists are presumably co-occurring or sharing similar ecological niches.

### Notable but less represented lineages

Although several lineages originally described for Crater Lake (with the prefix “CL”) (Urbach *et al*., 2001) were identified as representative hypolimnion specialists (Figs. 4–6), several other members dominant in Crater Lake were not highly represented in our study: CL120–10 of *Verrucomicrobia* (OTU65 in the present study; see Supplementary Data), CL0–1 of *Armatimonadetes* (OTU77), and CL500–9 of *Chloroflexi* (not detected in the present study). It is plausible that further investigations in lakes on other continents or those with depths > 250 m will detect bacterioplankton not found in the present study. Methanotrophs, such as *Methylococcaceae* and *Methylocystaceae*, also accounted for a very minor fraction of all amplicon reads (see Supplementary Data). A more intriguing result is the limited distribution and low relative abundance of LD12 (Figs. 2, 4), which is one of the most dominant and ubiquitous freshwater bacterioplankton (Zwart *et al*., 1998; Newton *et al*., 2011; Salcher *et al*., 2011). Notably, data produced using another sequencing platform (Miseq) from the same DNA samples showed a higher read proportion of LD12 (Fujinaga *et al*., unpublished data), despite the fact that both analysis used PCR primers that perfectly matched the LD12 16S rRNA sequence. Although the reason behind this discrepancy is unknown, it is possible that the relative abundance of LD12 was underestimated in the present study, which, in turn, might have overestimated other lineages among the reads, as in the discrepancy with the FISH results (Fig. S4). Direct cell enumeration using FISH should be considered an accurate abundance estimate.

### Potential ecotype diversification revealed by the oligotyping analysis

The oligotyping analysis revealed the intra-OTU diversification of the whole-layer generalists (Fig. 5). Some of the oligotypes were disproportionally distributed among depths or lakes (Fig. S2), suggesting that their ubiquity was achieved collectively by heterogeneous ecotypes that specialized in a respective niche. Such cryptic ecotype diversification within a ubiquitous freshwater lineage with an almost identical 16S rRNA sequence has previously been reported in *Limnohabitans* (Kasalický *et al*., 2013; Jezbera *et al*., 2013) and *Polynucleobacter* (Jezbera *et al*., 2011; Hahn *et al*., 2015, 2016), who reported diversification between habitats with different temperatures, pHs, organic and inorganic substrate availability, and geography. Recently, a horizontal oligotype profile in Lake Michigan indicated oligotype diversification within predominant bacterioplankton lineages between estuarine and pelagic sites (Newton and McLellan, 2015). The present study discovered micro-diversification between the epilimnion and hypolimnion, suggesting the presence of hypolimnion-specific ecotype within the common freshwater bacterioplankton. Further comparative studies of individual ecotypes will clarify their adaptation strategies to the respective water layers, which differ considerably in temperature, substrate availability, and grazing pressure.

In contrast to the whole-layer generalists, the low oligotype diversity of the hypolimnion specialists (Fig. 5) was intriguing, as we expected that the ecotypes would be diverse between the hypolimnia of different lakes, which are physically isolated and differ in physicochemical properties (Table 1). The occurrence of CL500–3 in TR (Fig. 6), which is a reservoir constructed just 10 years prior to the sampling, support the idea that hypolimnion specialists migrate between lakes; thus, diversification is limited. On the other hand, diverse ecotypes are likely present at least on a continental scale, as CL500–11 from European and North American lakes have a conserved single nucleotide difference from the sequences read in the present study (Fig. S7).

Contrary to the lineages described above, acI-C2 and LD19 were separated into epilimnion‐ and hypolimnion-specific OTUs by the > 97% OTU clustering threshold (Fig. 4), and the hypolimnetic OTUs were composed of an almost identical oligotype (Fig. 5). This indicates that the phylogenetic resolution required to resolve intra-OTU diversification within a given sequencing region depends on the lineage. As the present study used the finest phylogenetic resolution (i.e., single-nucleotide level) but analyzed only a 350 bp (V4 + V5 of the 16S rRNA gene) region, further micro-diversification studies should be carried out by increasing the number of sequenced regions.

### Conclusion

This study provides the first comprehensive overview of the bacterioplankton community inhabiting the oxygenated hypolimnion in 10 deep freshwater lakes with a variety of environmental characteristics. The community was composed of members occurring in common regardless of water layer or lake (whole-layer generalist) and members that were exclusively distributed in the hypolimnion (hypolimnion specialists). CARD-FISH revealed that predominant hypolimnion specialists, represented by *Chloroflexi*, *Planctomycetes*, and *Thaumarchaeota*, together accounted for 1.5–32.9% of the total bacterioplankton in the hypolimnion of the lakes. Moreover, an oligotyping analysis revealed the potential diversification of ecotypes in the common freshwater lineages (e.g., acI and *Limnohabitans*) among the lakes and water layers. These results reveal the uniqueness, ubiquity, and quantitative importance of bacterioplankton inhabiting the oxygenated hypolimnion. Collectively, the present results provide valuable suggestions for future ecological studies on deep freshwater ecosystems and eco-physiological and phylogeographical perspectives of general microbial ecology.

## ACKNOWLEDGMENTS

We are grateful to Y. Goda, T Denboh, Y Ito, M Sugiyama, and Japan Water Agency for their assistance in field sampling. This work is supported by JSPS KAKENHI Grant No. 15J00971 and 15J01065, and by the Environment Research and Technology Development Fund (No. 5-1304 and 5-1607) of the Ministry of the Environment, Japan.

Supplementary Information for this paper is available on the website.

